# A simple, ultrastable, and cost-effective oxygen-scavenging system for long-term DNA-PAINT imaging

**DOI:** 10.1101/2025.07.14.664726

**Authors:** Rebecca T. Perelman, George M. Church, Johannes Stein

**Affiliations:** Wyss Institute for Biologically Inspired Engineering, Harvard University, Boston, MA, USA; Harvard Biophysics Program, Harvard University, Boston, MA, USA; Department of Genetics, Harvard Medical School, Boston, MA, USA; Harvard-MIT Program in Health Sciences and Technology, Cambridge, MA, USA

## Abstract

DNA-PAINT (Points Accumulation in Nanoscale Topography) is a super-resolution microscopy technique capable of nanoscale imaging through the transient binding of fluorescently labeled imager strands to complementary DNA docking strands. Imager strands can be continuously replenished from an effectively infinite pool, making DNA-PAINT inherently resistant to photobleaching. However, extended DNA-PAINT imaging is limited by the formation of reactive oxygen species (ROS), which damage docking strands and reduce localization sampling over time. Although the state-of-the-art oxygen-scavenging system (OSS) can mitigate this damage, its enzymatic components degrade over time, reducing its performance, robustness and utility. Here, we introduce a simple, enzyme-free oxygen scavenging buffer based on sodium sulfite (Na_2_SO_3_) that overcomes these challenges. Our optimized formulation, combining Na_2_SO_3_ with Trolox (SST), effectively preserves docking strand integrity for over 24 hours and enhances long-term imaging performance. SST improves buffer stability tenfold, is easy to prepare and reduces costs by more than 90%, providing a robust, cost-effective, high-performance OSS for extended DNA-PAINT imaging.

## Main Text

Super-resolution microscopy has become a powerful tool for visualizing biological structures at spatial resolutions beyond the diffraction limit of light^[1–5]^. Single-molecule localization microscopy^[6]^ (SMLM) achieves sub-diffraction spatial resolution by temporally separating fluorescence emission through stochastic blinking. While highly effective, SMLM techniques can be limited by the irreversible loss of fluorescence due to photochemical damage during excitation, a phenomenon known as photobleaching. Photobleaching prevents repeated sampling of molecular locations and ultimately limits achievable resolution. DNA-PAINT, which achieves nanometer-precision localization through the transient binding of fluorescently labeled imager strands that produce stochastic blinking, offers resistance to photobleaching by continuously replenishing fresh imager strands from a large diffusing pool^[7,8]^. This reversible binding mechanism provides continuous resampling and theoretically unlimited localizations, enabling extended imaging^[9]^, molecular counting^[10,11]^, and resolution down to the level of single proteins^[12,13]^.

However, irreversible photo-induced damage to docking strands by reactive oxygen species (ROS) remains a limitation in DNA-PAINT, restricting imaging durations and affecting both achievable resolution and quantitative interpretations^[14]^. Oxygen scavenging systems (OSSs) are widely used in SMLM to mitigate photobleaching by removing molecular oxygen, thereby extending fluorophore lifetime^[6,14,15]^. In DNA-PAINT, OSSs also protect docking strands by limiting ROS generation^[14]^ (**Fig. 1a**). The state-of-the-art DNA-PAINT OSS^[8,16]^, denoted as ‘PPT’, contains protocatechuic acid (PCA), protocatechuate-3,4-dioxygenase (PCD), and the triplet state quencher Trolox. PPT maintains optimal photophysical conditions and effectively protects docking strands for up to ∼1-2 hours^[17]^. However, in the presence of oxygen, the PCD-driven reaction gradually acidifies the buffer (**Fig. 1b**, inset), reducing scavenging efficiency^[17]^ and altering DNA binding dynamics^[18]^. As a result, frequent buffer replacement is required, posing challenges for high-throughput or multiplexed DNA-PAINT workflows that require tens of sequential imaging rounds^[13,19,20]^ or for fluidics-based automation^[21]^. In addition, PPT preparation is labor-intensive and costly, limiting its scalability.

**Figure 1.**
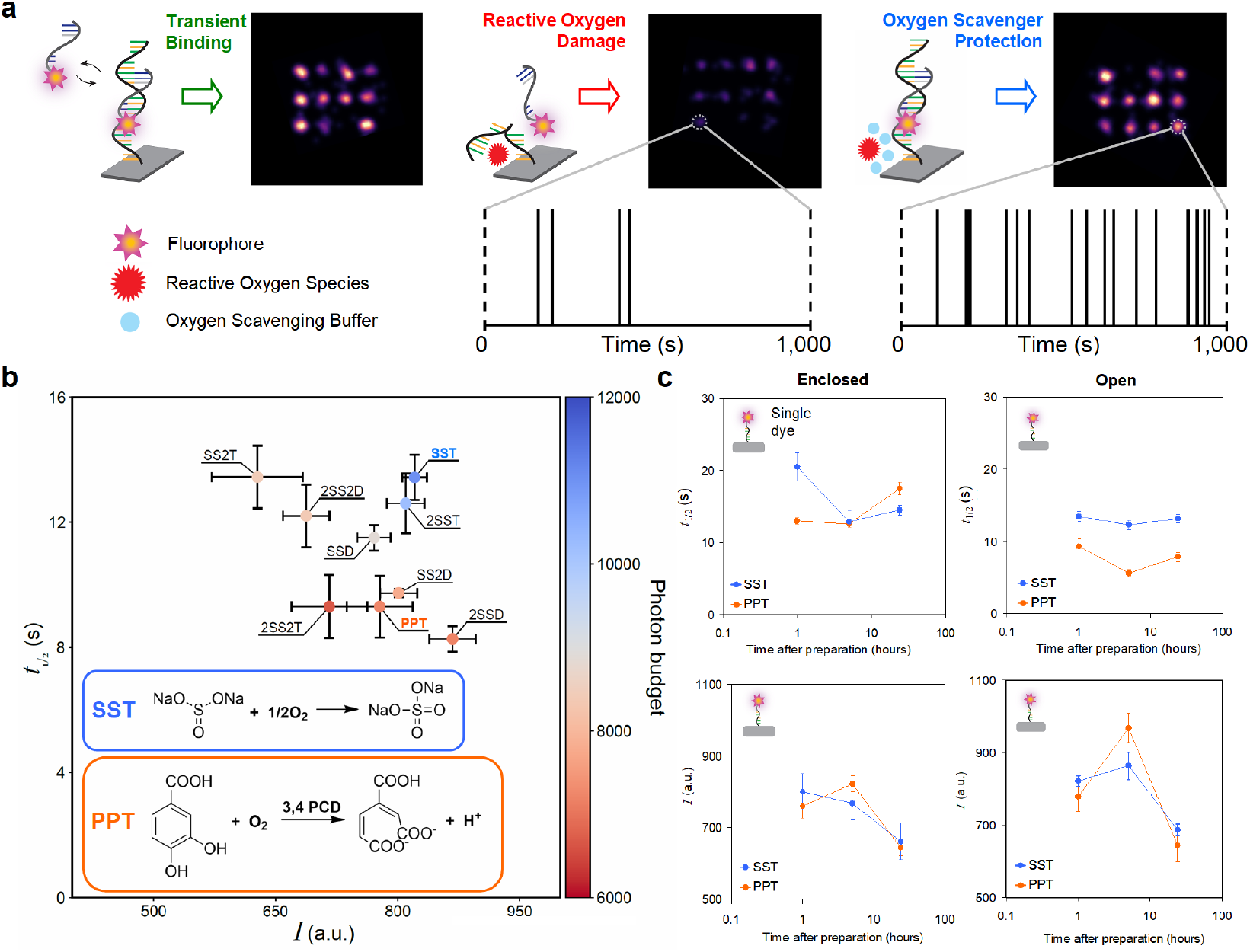
Stability and efficiency of DNA-PAINT oxygen-scavenging systems. **a**, Schematic illustration of transient imager-docking strand binding and ROS-induced damage. The protective effects of oxygen scavengers are demonstrated by the number of localizations in super-resolved images of a 20-nm DNA origami grid. In the absence of scavengers, ROS damage leads to incomplete sampling of docking strands. In their presence, docking strands remain fully accessible, yielding efficient localization sampling and continuous imager binding over time. **b**, Measured fluorescence half-life, *t*_1/2_, vs. average number of photons detected per localization, *I*. The corresponding photon budget (defined as *t*_1/2_ × *I*) is represented by the color scale indicated in the bar on the right. Conditions include SST, SSD, and PPT, as well as modified variants: 2SST and 2SSD (with doubled Na_2_SO_3_ concentrations), SS2T and SS2D (with doubled triplet state quenchers), and 2SS2T and 2SS2D (with both Na_2_SO_3_ and triplet state quencher doubled). Inset: oxygen scavenging mechanisms for Na_2_SO_3_ and PCD. **c**, Fluorescence half-life, *t*_1/2_ (top) and average number of photons detected per localization, *I* (bottom) measured as soon as possible after preparation and at 5 and 24 hours thereafter for PPT (orange) and SST (blue) using single-dye origami. Time after preparation is plotted on a logarithmic scale. Measurements were conducted in an enclosed chamber (left panel) and open chamber (right panel). Data are presented as means ±SEM (n=3).

Sodium sulfite (Na_2_SO_3_), a chemical oxygen scavenger (**Fig. 1**b, inset), has recently been applied in the context of fluorogenic DNA-PAINT^[22]^, where ROS formation is intrinsically reduced as quencher-labeled imager strands are protected from photobleaching during diffusion^[23]^. However, its potential as standalone OSS for conventional DNA-PAINT remains unexplored and a quantitative comparison with PPT is missing. Given its success in stabilizing imaging buffers for the SMLM variant STORM^[24–26]^ (Stochastic Optical Reconstruction Microscopy), Na_2_SO_3_ presents a promising, simpler, and more cost-effective alternative that warrants systematic assessment.

We systematically evaluated Na_2_SO_3_-based buffer formulations for their performance in DNA-PAINT imaging. We screened various Na_2_SO_3_ concentrations with the triplet-state quenchers Trolox or DABCO^[25,26]^ in single-dye experiments and identified Na_2_SO_3_ and Trolox (SST) as the optimal formulation for conventional DNA-PAINT. SST showed 1.4-to 2.2-fold greater photostability than PPT and preserved docking strand integrity over 24 hours, resulting in a tenfold improvement in buffer stability and ∼90% reduction in cost, with SST costing ∼$0.02/mL (primarily due to Trolox) compared to ∼$0.31/mL for PPT. SST enables long-term DNA-PAINT imaging by minimizing ROS formation and preserving docking strand integrity. It remains stable at room temperature for weeks, supports extended acquisitions without compromising image quality or the imager binding frequency, thereby improving the spatial resolution through enhanced sampling density^[6,27]^.

To evaluate Na_2_SO_3_ performance in combination with different triplet-state quenchers, we started with 30 mM Na_2_SO_3_, the optimal concentration identified for STORM imaging^[25]^, and formulated two buffers: one with 1 mM Trolox (SST), as used in PPT^[8]^, and the other with 65 mM DABCO^[25]^ (SSD). We also prepared variants with doubled concentrations of both Na_2_SO_3_ and triplet-state quenchers. Photostability was assessed using Cy3B-labeled single-dye DNA origami structures^[28,29]^, the most robust fluorophore for 560 nm excitation in DNA-PAINT^[30]^ (**Fig. S1**a). We measured the fluorescence half-life, *t*_1/2_, defined as the time at which half the dyes were photobleached upon continuous illumination, and the average number of photons detected per localization, *I*. Consistent with previous reports^[29]^, single Cy3b dyes bleached on the timescale of tens of seconds. SST showed the best performance, with a 40% in *t*_1/2_ (13.4 s vs. 9.3 s) and 50% in photon budget (∼12,000 vs. 8,000) compared to PPT (**Fig. 1**b).

Next, we evaluated SST performance and stability over 24 hours compared to PPT, using single-dye DNA origami in enclosed and open chambers to control oxygen exchange. In the enclosed chamber, which prevented oxygen influx, both buffers performed similarly (**Fig. 1**c, left). In the open chamber, where oxygen exchange was permitted and mimics typical DNA-PAINT conditions^[9]^, the *t_1/2_* of PPT declined over the 24-hour period, dropping by 40% within 5 hours of preparation, while SST showed only a 10% decline, with values comparable to its performance in the sealed chamber (**Fig. 1**c, right). After 5 hours, the *t_1/2_* in SST was more than twice that of PPT, demonstrating its superior photostability and long-term performance.

To assess whether SST minimizes photoinduced docking strand damage, we employed a DNA origami structure featuring twelve docking strands spaced 20 nm apart^[8]^ (**Fig. S1**b). As in previous work^[14]^, we conducted extended DNA-PAINT acquisitions of 25,000 frames at 200 ms exposure (∼1.5 h), during which freshly prepared PPT is known to preserve docking strands^[14]^. Since ∼5,000 frames are sufficient to localize all sites, we compared super-resolved images from the initial and final 5,000 frames (referred to as ‘first segment’ vs. ‘last segment’, hereafter) to quantify photoinduced damage and imaging fidelity^[14]^.

To evaluate the long-term stability of OSS performance, we conducted two rounds of extended DNA-PAINT acquisitions for each condition (PPT and SST): one immediately after OSS preparation and another 24 hours after OSS preparation. Between experiments, samples were stored at room temperature in their respective OSSs. All imaging conditions, including DNA origami and imager strand concentrations, as well as excitation laser power, were held constant. Super-resolved DNA origami images were reconstructed using identical parameters (**Fig. 2**a). Localization precision (*σ*_NeNA_) for each dataset was quantified using the nearest-neighbor analysis (NeNA) method^[31]^, which estimates the localization uncertainty at individual, spatially defined docking strands. Immediately after preparation, both OSSs yielded comparable image quality and *σ*_NeNA_ values in the first segment: 2.74 nm for SST and 2.71 nm for PPT; **Fig. 2**a, top left). After 1.5 hours of imaging, as expected, only minor signs of deterioration were observed in PPT, including reduced localization density and occasional docking strand loss in the last segment. In contrast, SST maintained image integrity without detectable deterioration (**Fig. 2**a, bottom left. See **Fig. S2** for additional 100 exemplary DNA origami per OSS).

**Figure 2.**
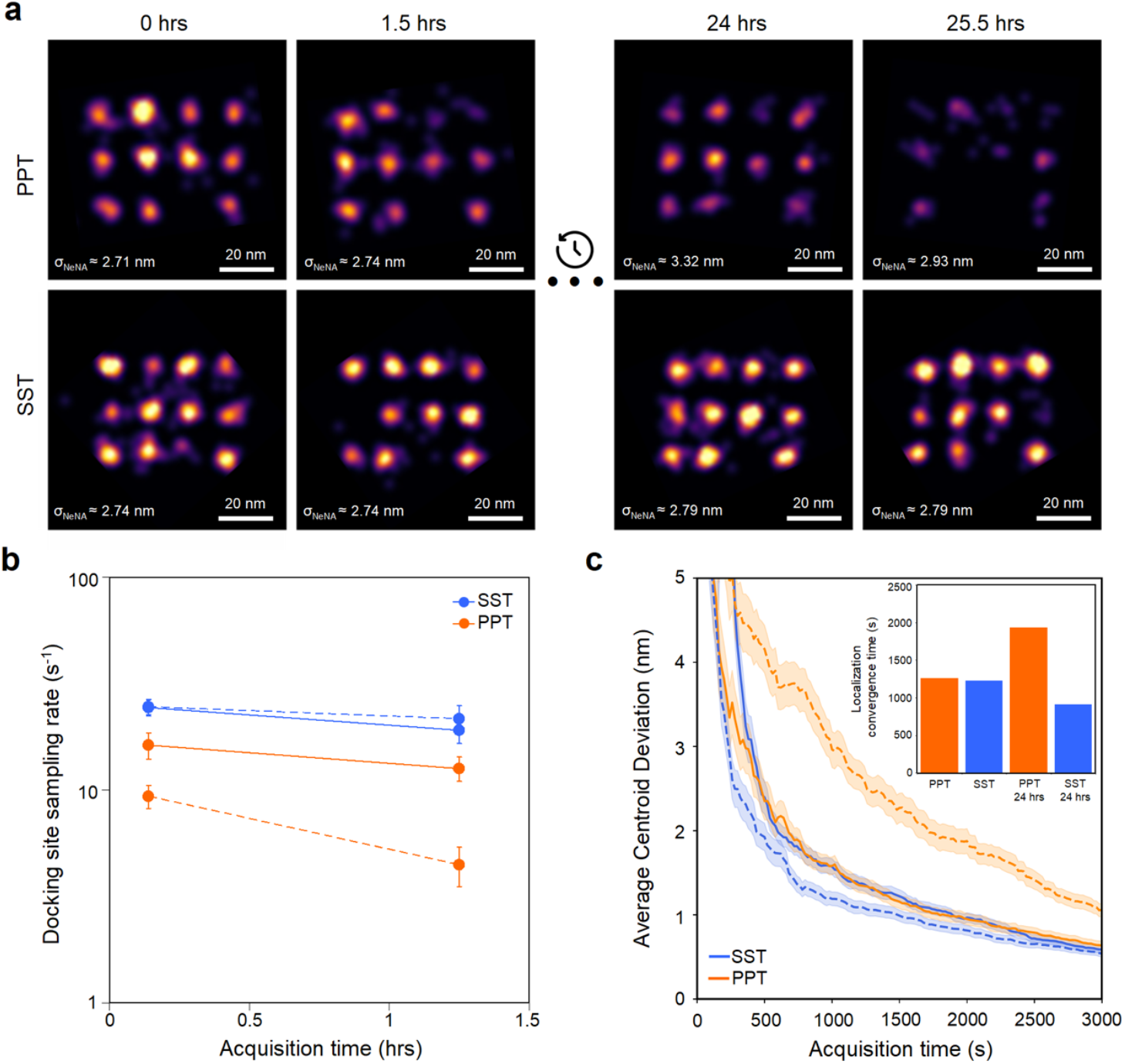
SST enhances DNA-PAINT docking strand preservation, resolution, and sampling density, as evaluated by extended imaging of DNA origami and analysis of damage rates. **a**, Representative DNA origami images acquired in PPT and SST (top and bottom, respectively) over 25.5 hours, including two 1.5-hour imaging sessions separated by 24 hours of idle time. In PPT, image intensity and quality progressively declined, whereas in the SST, they remained stable. The σ_NeNA_ localization precision is stated in each image. All imaging was performed at the same imager concentration, illumination (10 mW), magnification (100X), exposure time (200 ms), and images were rendered using the same parameters. Scale bars: 20 nm. **b**, Docking strand sampling rate in both buffers on logarithmic scale. While both buffers showed a similar decline in docking strand sampling rate during the first 1.5-hour imaging session (solid lines), the rate in PPT continued to decline over the 24-hour idle period, whereas it fully recovered in SST. In the second 1.5-hour session, the rate remained stable in SST but dropped significantly in PPT (dashed lines). Data are presented as means ±SEM (n=11). **c**, Convergence behavior of RESI center localizations relative to the full 25,000-frame dataset. Both buffers exhibited similar convergence rates immediately after preparation (solid lines), whereas SST showed markedly faster convergence than PPT after 24 hours (dashed lines). Data are presented as means ±SEM (n=100).

After 24 hours of storage, the performance differences between both OSSs became more pronounced. When extended DNA-PAINT acquisitions were performed at a new field of view, PPT exhibited a significant decline in image quality as early as in the first segment. This was evident in reduced localization sampling and increased localization uncertainty (*σ*_NeNA_ = 3.32 nm), and clear signs of damage to several docking strands. Over the following 1.5 hours, degradation continued, and by the last segment, many docking strands were barely detectable (**Fig. 2**a, top right). In striking contrast, SST preserved high localization precision (∼2.79 nm) and dense localization sampling throughout the entire acquisition, with minimal apparent docking strand damage even after prolonged imaging (**Fig. 2**a, bottom right. See **Fig. S3** for additional 100 exemplary DNA origami per OSS).

To further quantify these differences, we measured the number of binding events per docking strand, defined as the docking strand sampling rate (s^-1^), in both the first and last segments of the datasets immediately after buffer preparation and again after 24 hours. To ensure reproducibility, sampling rate measurements were replicated across three independently prepared samples (**Fig. S4**a). Notably, SST supported nearly a twofold higher docking strand sampling rate compared to PPT in the initial segments (∼17 s^-1^ vs. ∼9 s^-1^), despite identical imager strand concentrations. This suggests that SST not only preserves docking strand integrity over time, but also substantially enhances hybridization kinetics.

Following preparation, both OSSs exhibited a comparable reduction in active docking strands between the first and last segments (20% decrease in SST vs. 23% in PPT). However, after 24 hours of storage, docking strand activity continued to decline in PPT, whereas it fully recovered in SST, consistent with trends observed in **Fig. 2**a. During the second 1.5-hour imaging session, the docking strand sampling rate remained stable in SST (13% decrease), while a marked decline was measured in PPT (52% decrease) (**Fig. 2**b). We further evaluated both OSSs one month after preparation. Strikingly, SST maintained high performance with only a 10% decrease in sampling rate over the 1.5-hour imaging round, while PPT showed substantial degradation, marked by a 54% decrease (**Fig. S4**b). These findings suggest that preparing larger buffer stocks for extended use may be advantageous, although we note that the one-month storage experiment was performed only once.

Next, we examined the effects of each OSS on additional technical aspects of DNA-PAINT imaging. A recent advancement, RESI (Resolution Enhancement by Sequential Imaging), introduced by Reinhardt et al., offers a powerful approach to enhance localization precision^[13]^. RESI builds on the concept of performing Exchange-PAINT^[9]^ on a single target species to generate sparse localization clouds, each corresponding to an individual molecule. By precisely determining the center of these ‘single-docking strand’ clouds, the method enables pinpointing docking strand positions with reported precision down to the Ångström scale^[13]^. Due to their sequential nature, both RESI and Exchange-PAINT require docking strands to remain intact over extended imaging times, exceeding those typical of conventional DNA-PAINT. In this context, we sought to assess the compatibility of SST with such demanding imaging protocols.

Given that our DNA origami design enables single docking strand resolution, we leveraged our extended DNA-PAINT datasets to assess performance in the context of RESI analysis. Briefly, RESI localizations were calculated by determining the weighted mean of the x- and y-coordinates from all localizations within a given localization cloud. For each of the four conditions (PPT fresh, PPT after 24 hours, SST fresh, and SST after 24 hours) we randomly selected 100 individual docking strands and computed their RESI center localizations from the full 25,000-frame dataset, serving as our ground-truth reference. To evaluate convergence behavior, we analyzed subsets of the data at shorter acquisition durations, quantified the deviation of each RESI localization from the full-data estimate, and determined the imaging time required for RESI localizations to converge to that reference. As expected, both SST and PPT performed comparably immediately after preparation, requiring approximately 21 minutes of imaging to achieve an e-fold improvement in localization precision. However, after 24 hours, RESI convergence in PPT was markedly impaired, requiring up to 60% longer acquisition times to achieve comparable accuracy, highlighting a clear decline in performance. In contrast, SST not only preserved but even improved its performance after 24 hours, reaching the same precision threshold in just 15 minutes compared to 21 minutes previously, further underscoring its suitability for RESI applications (**Fig. 2**c). These results demonstrate that SST not only protects docking strands from damage but also preserves consistent localization sampling, enabling robust high-precision imaging even over extended acquisition times.

For unknown biological structures, the positions of docking strands are not known in advance. Therefore, achieving high a localization density is essential to ensure that structural features are adequately resolved. According to the Nyquist-Shannon sampling criterion^[32]^, resolution is fundamentally limited to twice the average spacing between localizations, meaning that a reduction in available docking strands over time directly constrains the highest resolvable spatial frequencies^[27]^. In **Supplementary Note 1** we discuss the theoretical implications on resolution loss on unknown structures based on the docking strand damage rates obtained from DNA origami experiments (**Fig. S4**c,d).

While DNA origami serves as an excellent benchmark for assessing docking strand damage in DNA-PAINT experiments, it does not fully reflect the complexity of cellular imaging. To assess the suitability of SST for cellular experiments and to compare overall image quality, we performed extended DNA-PAINT acquisitions targeting microtubule filaments in fixed HeLa cells. Imaging was carried out using both PPT and SST, immediately after buffer preparation and again after a 24-hour storage period. As before, each acquisition comprised 25,000 frames, allowing comparison between the initial and final 5,000-frame segments. **Fig. 3**a shows the super-resolved microtubule reconstructions for both OSSs over the full acquisition period immediately after preparation and demonstrates that SST performs at least as well as PPT in cellular imaging applications. Notably, a substantially lower imager concentration was used in the microtubule experiments compared to the DNA origami assays. Since ROS-induced docking strand damage is concentration-dependent^[14]^, the performance difference between SST and PPT was less pronounced in microtubule imaging. Nevertheless, a significant increase in docking site localization density was observed in SST over PPT (**Fig. 3** and **S5**a).

**Figure 3.**
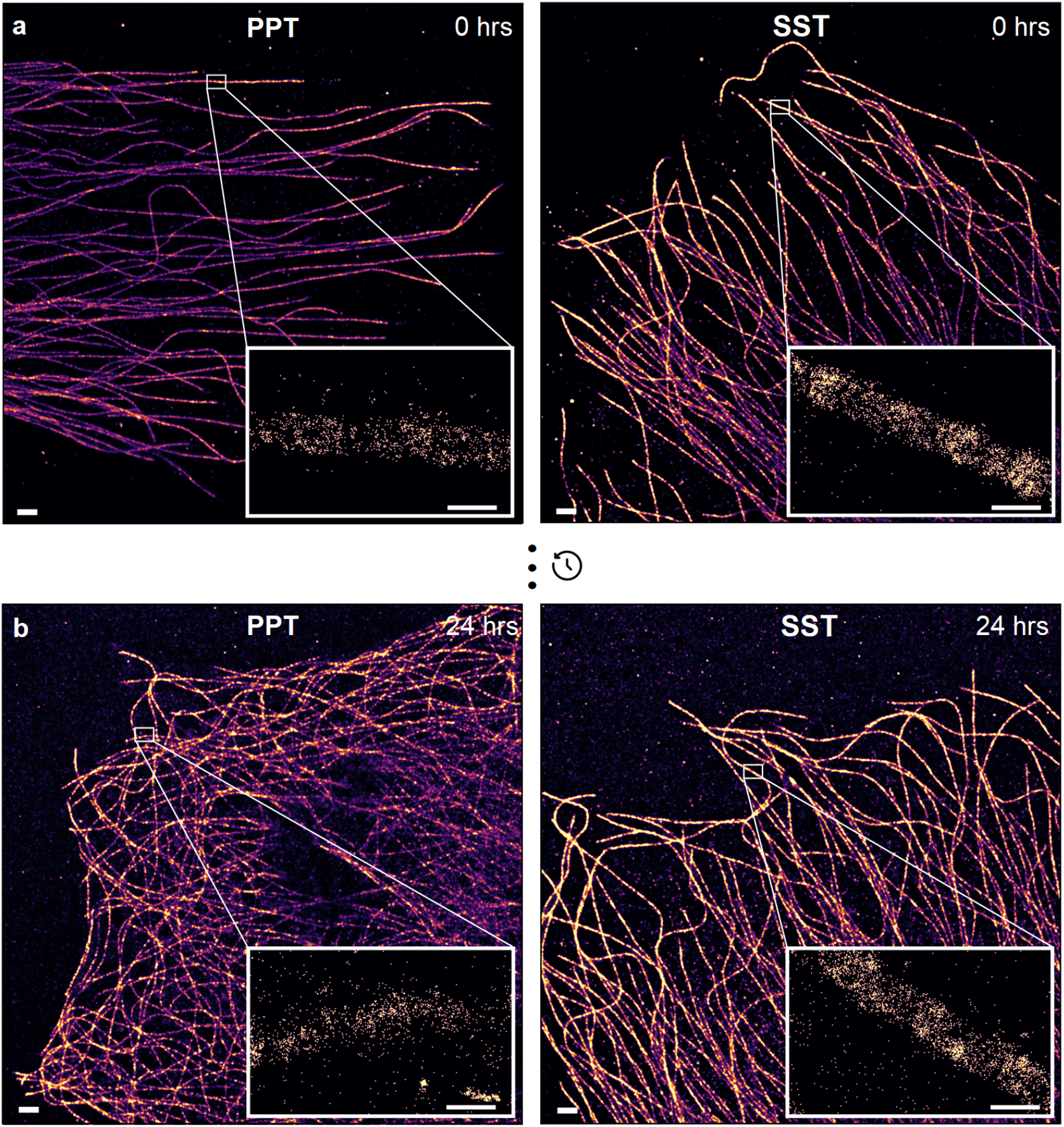
The advantages of SST extend to DNA-PAINT imaging in fixed cells. **a**, DNA-PAINT images of the microtubule network in fixed HeLa cells acquired in PPT (left) and SST (right). Images were acquired at the same imager concentration, illumination (10 mW), magnification (100X), exposure time (200 ms), and displayed using the same intensity range. Inset: zoom-in views range rendered at the same parameters. **b**, DNA-PAINT images of the microtubule network in fixed HeLa cells acquired in PPT (left) and SST (right) 24 hours after preparation. Images were acquired under the same conditions. Inset: zoom-in views rendered at the same parameters. Scale bars: 1000 nm. Scale bars: 100 nm. FOV: 130 × 130 μm^2^.

This effect was further quantified by measuring the docking strand sampling rate across microtubule segments (**Fig. S**5b). PPT showed a ∼30% decline between sessions, compared to ∼10% in SST. Moreover, SST yielded over twice the localization density of PPT, highlighting its improved retention of docking strand activity and imaging performance. To assess structural preservation, cross-sectional fluorescence intensity profiles of microtubules imaged in PPT and SST at 0 and 24 hours were analyzed. All profiles reflected the expected hollow, rod-like morphology with peak-to-peak distances of ∼38 nm, consistent with reported values^[33,34]^ (**Fig. S**6). These findings demonstrate that SST is fully compatible with cellular DNA-PAINT imaging and that its in vitro performance improvements translate robustly to cellular contexts.

In summary, we present SST as an ultra-stable, and highly efficient oxygen-scavenging system for conventional DNA-PAINT imaging that significantly enhances image quality, extends imaging durations, and improves resolution by increasing photostability and minimizing docking strand damage. While alternative strategies exist for reducing molecular oxygen, including vacuum systems, DNA-mediated delivery of photostabilizers^[35]^, and fluorogenic dyes that resist photobleaching^[22]^, these approaches tend to introduce additional complexity and cost. In contrast, SST is simple, cost-effective, and easy to implement using standard laboratory reagents.

The advantages of SST demonstrated here are based on its compatibility with Cy3B, a high-performance orange fluorophore widely used in DNA-PAINT^[8,13,19,20,30]^. Recently introduced ‘ANice’, a non-toxic and cost-effective OSS optimized for green fluorophores^[36]^, reflects a broader push toward broadly-accessible imaging buffers. Although no universal OSS exists, comprehensive fluorophore screens provide a foundation for adapting SST to additional dyes^[30]^. This is particularly important for multicolor DNA-PAINT applications requiring spectral demixing^[37]^.

Beyond DNA-PAINT, a major advantage of SST lies in its ability to preserve DNA docking strands during prolonged imaging sessions – a critical requirement for a wide range of DNA-based multiplexed techniques, such as Oligopaints/OligoSTORM^[38,39]^, MERFISH^[40]^, seqFISH^[41]^, SABER^[42]^, or OligoFISSEQ^[43]^, which rely on fixed DNA labels and barcoding strategies. By minimizing oxidative damage and maintaining the stability of DNA labels, SST has the potential to enhance the robustness and fidelity of these imaging strategies.

Together, our findings position SST as a versatile and accessible enhancement to DNA-PAINT and as a strong foundation for DNA-based multiplexed imaging.

## Supporting Information

The methods and chemicals for the design and preparation of our samples are described in the SI Materials and Methods section. Instrumentation and procedures for data acquisition as well as for data analysis are described in the SI Materials and Methods section.

Supplementary PDF containing the materials and methods sections, Supplementary Figures and Supplementary Tables.

The authors have cited additional references within the Supporting Information.^[44]^

## Author contributions

R.T.P. performed experiments, analyzed data and wrote the manuscript. J.S. analyzed data and wrote the manuscript. J.S. conceived the study and J.S. and G.M.C. supervised the study. All authors read and approved the manuscript.

## Funding

R.T.P acknowledges support by the National Science Foundation Graduate Research Fellowship Program. G.M.C. acknowledges support by the Department of Energy (DE-FG02-02ER63445). This work was also supported by an NIH award to Chao-ting Wu (5RM1HG011016-03). J.S. acknowledges support by the European Molecular Biology Organization (ALTF 816-2021).

## Acknowledgements

We thank Ryan McMillan, Michel Nofal and Maurice Perez for help with cell culture. We thank Michel Nofal and Ryan McMillan for support with DNA-antibody conjugation and helpful discussions. We thank Jenny Tam, Tom Ferrante, Maurice Perez and Jennifer Shay for providing infrastructure and experimental support.

## Conflict of interest

Potential conflicts of interest for G.M.C. are listed on https://arep.med.harvard.edu/t/. All other authors declare no competing financial interest.

---

**Table of Contents.**
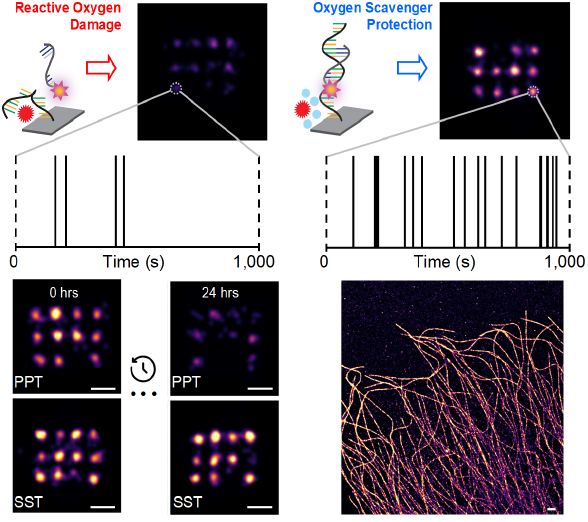
A simple, enzyme-free, and cost-effective oxygen scavenging system enhances DNA-PAINT super-resolution microscopy by increasing photostability, minimizing docking strand degradation, and enabling faster, high-precision localizations. Its long-term stability supports extended imaging, yielding improved spatial resolution and image quality. This robust system provides a foundation for advancing a wide range of DNA-based multiplexed imaging techniques.

