## Supplementary Information for "A simple, ultrastable, and cost-effective oxygen-scavenging system for long-term DNA-PAINT imaging"

**Methods**

**Supplementary Figures**

**Supplementary Notes**

**Supplementary Tables**

**References**

### Methods

**Materials.** Unmodified, dye-labeled, and modified DNA oligonucleotides were purchased from Integrated DNA Technologies, Metabion and Biomers. Unmodified oligos were purified via standard desalting and modified oligos via HPLC. DNA scaffold strands were purchased from Tilibit (p7249, identical to M13mp18). Sample chambers were ordered from Ibidi GmbH (8-well 80827 and 18-well 81817). Tris 1M pH 8.0 (AM9856), EDTA 0.5M pH 8.0 (AM9261), Magnesium 1M (AM9530G) and Sodium Chloride 5M (AM9759) were ordered from Ambion. Streptavidin (S-888) Ultrapure water (15568025), PBS (20012050), 4',6-Diamidino-2-Phenylindole, Dihydrochloride (D1306) (A39255), BSA (AM2616), DMEM (10569) and Dithiothreitol (DTT) were purchased from Thermo Fisher Scientific. BSA-Biotin (A8549), Tween-20 (P9416-50ML), Glycerol (cat. 65516-500ml), (+-)-6-Hydroxy-2,5,7,8-tetra-methylchromane-2-carboxylic acid (Trolox) (238813-5G), methanol (32213-2.5L), 3,4-dihydroxybenzoic acid (PCA) (37580-25G-F), protocatechuate 3,4-dioxygenase pseudomonas (PCD) (P8279-25UN), sodium sulfite (S0505-250G), 1,4-Diazabicyclo[2.2.2]octane (DABCO) (D27802-25G), Triton-X 100 (93443), Sodium Azide (S2002), and HEPES (H4034-100G) was purchased from Sigma-Aldrich. 10% fetal bovine serum was purchased from Genesee Scientific (25-514). EM grade gluyaraldehyde was purchased from Electron Microscopy Services (16220). 90 nm gold nanoparticles (G-90-20-10 OD10) were purchased from Cytodiagnostics. Primary anti a-tubulin (rabbit, 2125BF) antibody was purchased from Cell Signaling. Secondary goat anti-mouse labeled with Alexa488 antibody (A11029) was purchased from Thermo Fisher Scientific. 0.5-mL Amino Ultra Centrifugal Filters with 50 kDa and 10 kDa molecular weight cutoffs were purchased from Millipore (UFC5050 and UFC5010, respectively). DBCO-sulfo-NHS ester cross-linker was purchased from Vector Laboratories (CCT-A124). Qubit Protein Assay (Q33211), NuPage 4-12% Bis-Tris protein gels (NP0323BOX), NuPage LDS Sample Buffer (NP0007) was purchased from Invitrogen. InstantBlue Coomassie Protein Stain was purchased from Abcam (ab119211).

**Buffers.** Four buffers were used for sample preparation and imaging: Buffer A (10 mM Tris-HCl pH 7.5, 100 mM NaCl); Buffer B (5 mM Tris-HCl pH 8.0, 10 mM MgCl2, 1 mM EDTA); Buffer C (1× PBS, 500 mM NaCl); 10x folding buffer (100 mM Tris,10 mM EDTA pH 8.0, 125 mM MgCl2). Antibody storage buffer: 1% BSA, 0.1% Sodium Azide, 10 mM EDTA, 50% glycerol). Buffers were checked for pH. For imaging, Buffer C was supplemented with oxygen scavenging & triplet state quenching system PPT (1× PCA, 1× PCD, 1× Trolox) or SST (30 mM Sodium Sulfite, 1× Trolox) prior to imaging.

**PCA, PCD, Trolox.** 100× Trolox: 100 mg Trolox, 430 µL 100% Methanol, 345 µL 1 M NaOH in 3.2 mL H2O. 40× PCA: 154 mg PCA was mixed with 10 mL water adjusted to pH 9.0 with NaOH. 100× PCD: 9.3 mg PCD, 13.3 mL of buffer (100 mM Tris-HCl pH 8, 50 mM KCl, 1 mM EDTA, 50% glycerol).

**Sodium Sulfite, Trolox.** 100× Trolox: 100 mg Trolox, 430 µL 100% Methanol, 345 µL 1 M NaOH in 3.2 mL H2O. 1 M Sodium Sulfite: 1,764 mg Sodium Sulfite, 14 mL H2O, and can be left at room temperature for at least 1 month.

**DNA origami design and assembly.** DNA origami structures were designed using the Picasso Design[1] module. A list of all used DNA strands and their respective ordering conditions can be found in Supplementary Tables 1-3. We used two previously published DNA origami designs[1–3]: single dye (SD) and 20-nm grid origami structures (Fig. S1). The single dye origami structure had a single extension at the top side at position 2B07 of Picasso Design labeled permanently with a Cy3B molecule. The 20 nm grid was a 3×4 grid motif with 20-nm spaced docking strands. Folding of structures was performed using the following components: single-stranded DNA scaffold (0.01 µM), core staples (0.1 µM), biotin staples (0.01 µM), extended staples for DNA-PAINT (each 1 µM), 1x folding buffer in a total of 50 µl for each sample. Annealing was done by cooling the mixture from 80 °C to 25 °C in 3 hours in a thermocycler. Using a 1:1 ratio between scaffold and biotin staples allows sample preparation without prior DNA origami purification, where otherwise free biotinylated staples would saturate the streptavidin surface and prevent origami immobilization on the glass surface. As docking strand sequence, we used a 17 nt motif (TT 5×CTC), and as imager strand sequence we used a 7 nt motif (Pm2-Cy3B), which directly hybridized to the docking strand sequence.

**DNA origami sample preparation.** For enclosed chamber preparation, a flow chamber with an inner volume of 20 µL was formed as described in ref. [1] using a plasma cleaned coverslip (no. 1.5, 18 × 18 mm$^2$, 0.17 mm thick) placed on a plasma cleaned glass slide (3 × 1 inch$^2$ 1 mm thick) using two strips of double-sided tape (Scotch, cat. no. 665D). First, 20 µL of biotin-labeled bovine albumin (1 mg/ml, dissolved in Buffer A) was flown into the chamber and incubated for 2 min. Then the chamber was washed using 40 µl of Buffer A. Second, 20 µl of streptavidin (0.5 mg/ml, dissolved in Buffer A) was then flown through the chamber and incubated for 2 min. Next, the chamber was washed with 40 µl of Buffer A and subsequently with 40 µl of Buffer B. Then, 20 µl of single dye origami structures (1:100-200 dilution in buffer B from folded stock) were flushed into the chamber and incubated for 5 minutes. Afterwards the chamber was washed with 40 µl of Buffer B again. Finally, the Buffer C with the oxygen scavenging & triplet state quenching systems was flown into the chamber.

Ibidi 8-well slides were prepared as follows. A 10 µl drop of biotin582 labeled bovine albumin (1 mg/ml, dissolved in buffer A) was placed at the chamber center and incubated for 2 min and aspirated. The chamber was then washed with 200 µl of buffer A, aspirated, and then a 10 µl drop streptavidin (0.5 mg/ml, dissolved in buffer A) was placed at the chamber center and incubated for 2 min. After aspirating and washing with 200 µl of buffer A and subsequently with 200 µl of buffer B, a 10 µl of DNA origami (1:100-200 dilution in buffer B from folded stock) was placed at the chamber center and incubated for 5 min. Next, the chamber was washed 2× with 200 µl of Buffer B. Finally, Buffer C with the oxygen scavenging & triplet state quenching systems and imager strand (final concentration of ~235 pM) was added for DNA-PAINT imaging.

**Conjugation of secondary antibodies with docking strands.** DNA antibody conjugations were performed as previously described[4] in 0.5-mL Amino Ultra Centrifugal Filters with 50 kDa molecular weight cutoffs with DBCO-sulfo-NHS ester cross-linker, which was dissolved at 20 mM DMSO and stored in single-use aliquots at -80° C. This cross-linker links azide-functionalized DNA oligonucleotides to surface-exposed lysine residues. Azide-functionalized DNA oligonucleotides were stored in 1 mM deionized water. Critically, all antibodies were ordered carrier-free, as common preservatives such as bovine serum albumin and sodium azide interfere with the conjugation reaction. First, 500 µL PBS was added to the Amicon filters, which were centrifuged for 5 min at 10,000 rcf. After wetting the filters, 25 µg antibody was added and washed twice with PBS. For each wash, PBS was added to a total volume of 500 µL, and the filters were centrifuged for 5 min at 10,000 rcf. If after the second spin, the total volume remaining in each filter was greater than 100 µL, the filters were centrifuged again for 5 min at 10,000 rcf. After the second PBS wash, a 20-fold molar excess of DBCO-sulfo-NHS ester cross-linker and a 20-fold molar excess of DNA oligonucleotide were added, and after gentle mixing, each conjugation reaction was incubated in the dark at 4° C overnight. The following day, conjugated antibodies were washed three times with PBS, as described above. To elute the antibody, the filter was inverted in a fresh tube and centrifuged for 2 min at 1,500 rcf. The conjugated antibody was transferred to a clean tube and stored at -20° C in antibody storage buffer. Concentrations were measured using the Qubit Protein Assay. DNA-antibody conjugation was confirmed by comparing unconjugated and conjugated antibodies on NuPage 4-12% Bis-Tris protein gels. For each sample, 0.5 µg total protein was added to NuPage LDS Sample Buffer and 50 mM DTT. Protein was denatured at 80° C for 10 min. Gels were run at 75 V for 5 min, then at 180 V for 60 min. Gels were stained with InstantBlue Coomassie Protein Stain for 15 minutes at room temperature, rinsed with water, and imaged on a Sapphire Biomolecular Imager (Azure Biosystems).

**Cell culture and plating.** HeLa cells were maintained in DMEM supplemented with 10% fetal bovine serum at 37 °C with 5% $CO_2$ and were checked regularly for mycoplasma contamination. For imaging of whole HeLa cells, ~8K cells were seeded in each well of an Ibidi 18-well chamber, placed in the incubator overnight and fixed the following day.

**Fixation and labeling for HeLa cell imaging.** 24h after seeding HeLa cells in Ibidi 18-well chambers, cells were pre-fixed in 0.2% GA, 0.25% Triton X-100 in PBS for 90 seconds and then, fixed in 2% GA in PBS for 10 minutes at 37°C. Next, samples were washed 3× in PBS and then, quenched in 0.1% $NaBH_4$ in PBS for 7 minutes. Samples were then washed 4× in PBS (30s, 60s, 2×5 min), and both blocked and permeabilized in 3% BSA and 0.25% Triton X-100 in PBS at room temperature for 90 minutes. Primary rabbit anti-Pol II S5p antibody was added at 1:100 in 3% BSA and 0.1% Triton-X 100 in PBS and incubated overnight at 4°C. The next morning, samples were washed 4× washes in PBS (30s, 60s, 2× 5min) and DNA-conjugated secondary antibody (1:100) was added at 1:100 in 3% BSA and 0.1% Triton-X 100 in PBCS and incubated for 1h at room temperature. Samples were quickly washed 3× in PBS, incubated with gold particles as fiducial markers (1:20 in PBS) for 5 min, washed again 3× 5 min in PBS before adding Buffer C with the oxygen scavenging & triplet state quenching systems and imager (final concentration of ~4 pM) for DNA-PAINT imaging.

**Super-resolution microscopy setup.** TIRF and HILO imaging was carried out on a Nikon Ti inverted microscope equipped with a Nikon Ti-TIRF-EM Motorized Illuminator, a Nikon LUN-F Laser Launch with single fiber output (561 nm, 70mW) and a Lumencore SpectraX LED Illumination unit. The objective-type TIRF system with an oil immersion objective (Apo TIRF 100×/1.49 DIC N2). DNA-PAINT experiments were performed using the 560 nm laser line and fluorescence emission was passed through a Chroma ZT 405/488/561/640 multi-band pass dichroic mirror mounted on a Nikon TIRF filter cube located in the filter cube turret and a Chroma ET 595/50m band pass emission filter located on a Sutter emission filter wheel within the infinity space of the stand before image recording on a line on a sCMOS camera (Andor, Zyla 4.2) mounted to a standard Nikon camera port.

**Imaging conditions.** All fluorescence microscopy data was recorded with the sCMOS camera (2048 × 2048 pixels, pixel size: 6.5 µm). Both microscope and camera were operated with the Nikon Elements software at 2×2 binning and 1024 × 1024 pixel field-of-view. The camera read out rate was set to 200 MHz and the dynamic range to 16 bit. Laser power for imaging single dye origami structures was 0.1%, and 20-nm grids and microtubules was 35% (10 mW).

**Image analysis.** All DNA-PAINT imaging data was processed and reconstructed using the Picasso software suite. Briefly, a standard single molecule localization algorithm was applied to the raw SD and 20 nm grid origami image stacks to generate a pointillist super-resolution reconstruction[1]. For all surface-immobilized SD origami experiments, only fluorescent spots that appeared within the first five frames were localized. When measuring signal duration, interruptions of up to two frames were permitted without ending the trace. Following localization, the 20 nm grid DNA origami structures appeared as clusters of localizations after image correlation-based drift correction[1]. Several clusters were manually selected using a circular region with a diameter of one camera pixel, and similar clusters were automatically identified. This process was then repeated at higher resolution for individual docking strand localization clusters in the 20 nm grid, using a circular region with a diameter of 0.125 camera pixels.

Kinetic analysis was based on a custom Python package picasso_addon (https://github.com/schwille-paint/picasso_addon). For SD origami experiments, kinetic paremeters for each pick (e.g. number of localizations, photon counts, $t_{1/2}$ etc.) were obtained using the spt.immobile_props.main() function. For 20 nm grid origami and microtubule experiments, the docking site sampling rate was determined by counting localizations at each docking strand or within a defined region, respectively, over a specified time interval. Detailed information about the picasso_addon API is available at https://picasso-addon.readthedocs.io/en/latest/index.html for detailed information about.

### Supplementary Figures

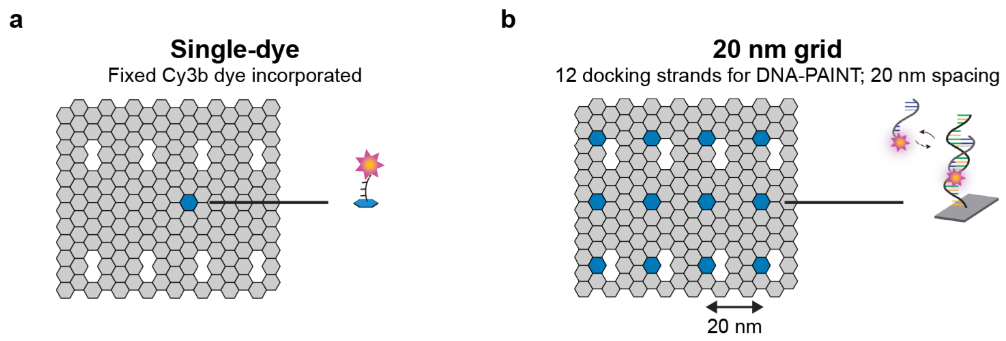


**Fig. S1 | DNA origami designs. a**, Single-dye (SD) DNA origami with fixed Cy3B. **b**, 3×4 20 nm DNA origami grid.

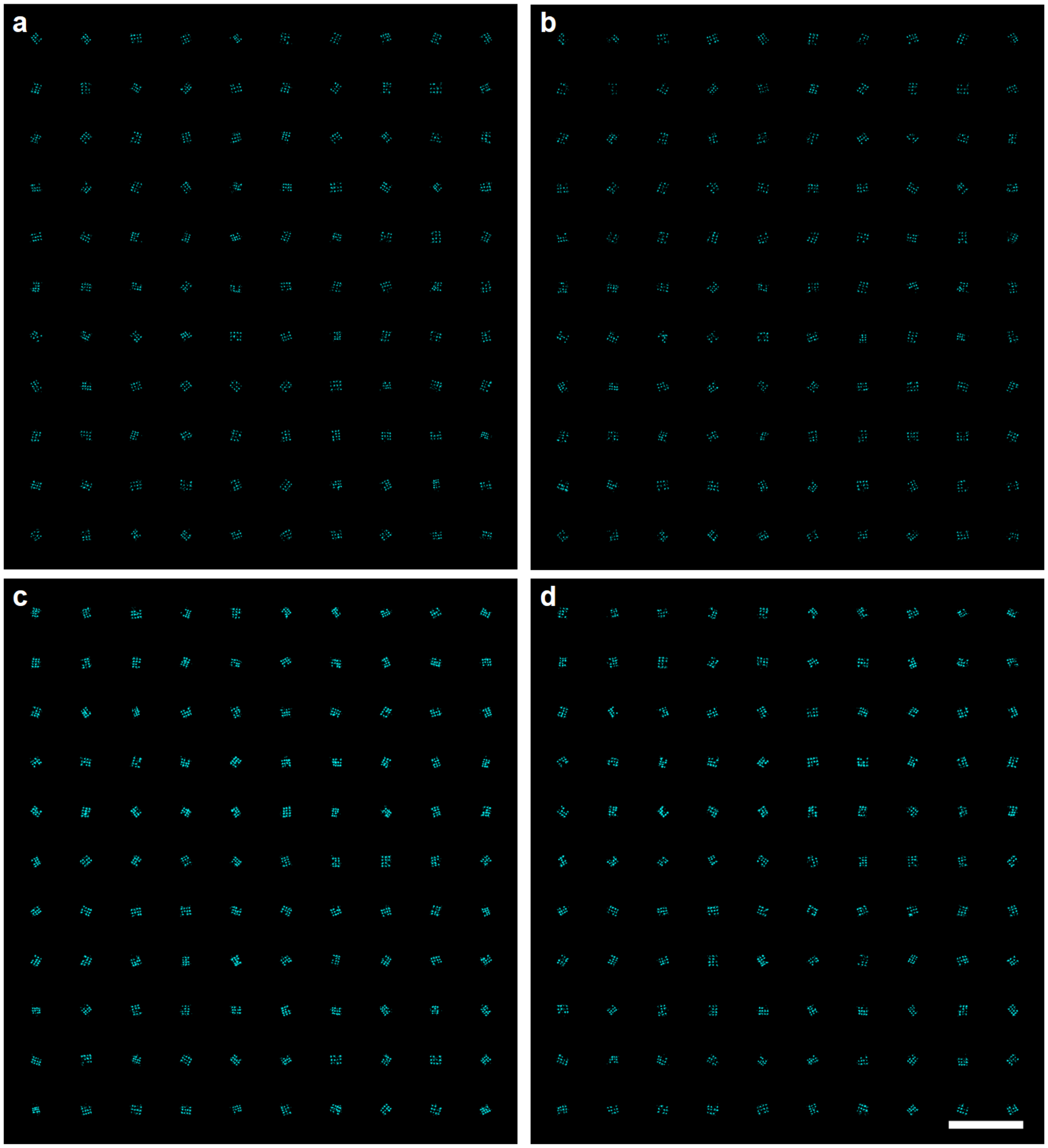


**Fig. S2 | Individual origami images in SST and PPT immediately after preparation. a**, Random selection of 110 origami structures in PPT over the first 5,000 frames, displayed in 10×11 grid. **b**, The same 110 origami structures imaged in PPT during the last 5,000 frames. **c**, Random selection of 110 origami structures in SST over the first 5,000 frames, displayed in 10×11 grid. **d**, The same 110 origami structures imaged in SST during the last 5,000 frames. Scale bar: 500 nm.

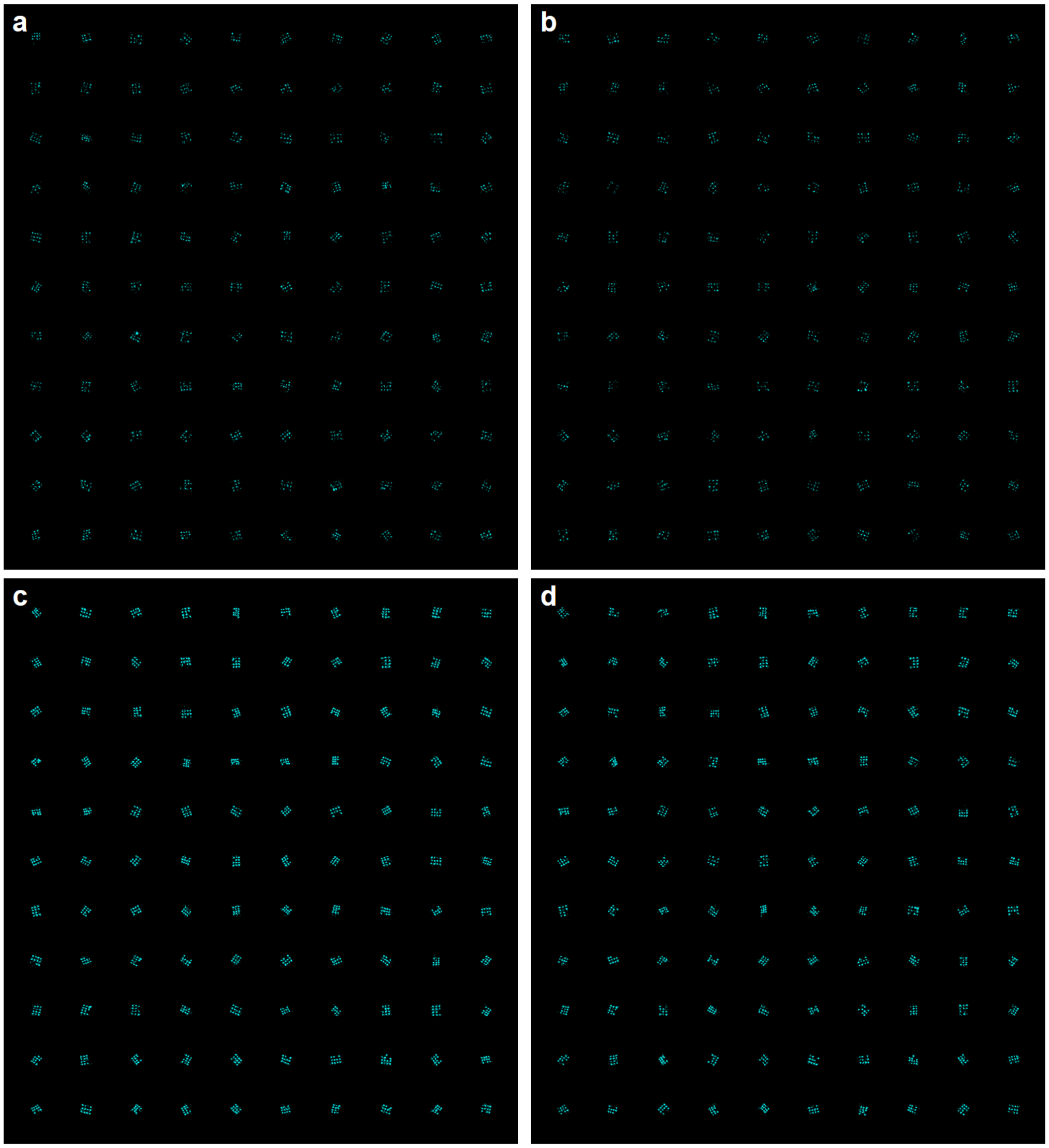


**Fig. S3 | Individual origami images in SST and PPT 24 hours after preparation.** **a**, Random selection of 110 origami structures in PPT over the first 5,000 frames, displayed in 10×11 grid. **b**, The same 110 origami structures imaged in PPT during the last 5,000 frames. **c**, Random selection of 110 origami structures in SST over the first 5,000 frames, displayed in 10×11 grid. **d**, The same 110 origami structures imaged in SST during the last 5,000 frames. Scale bar: 500 nm.

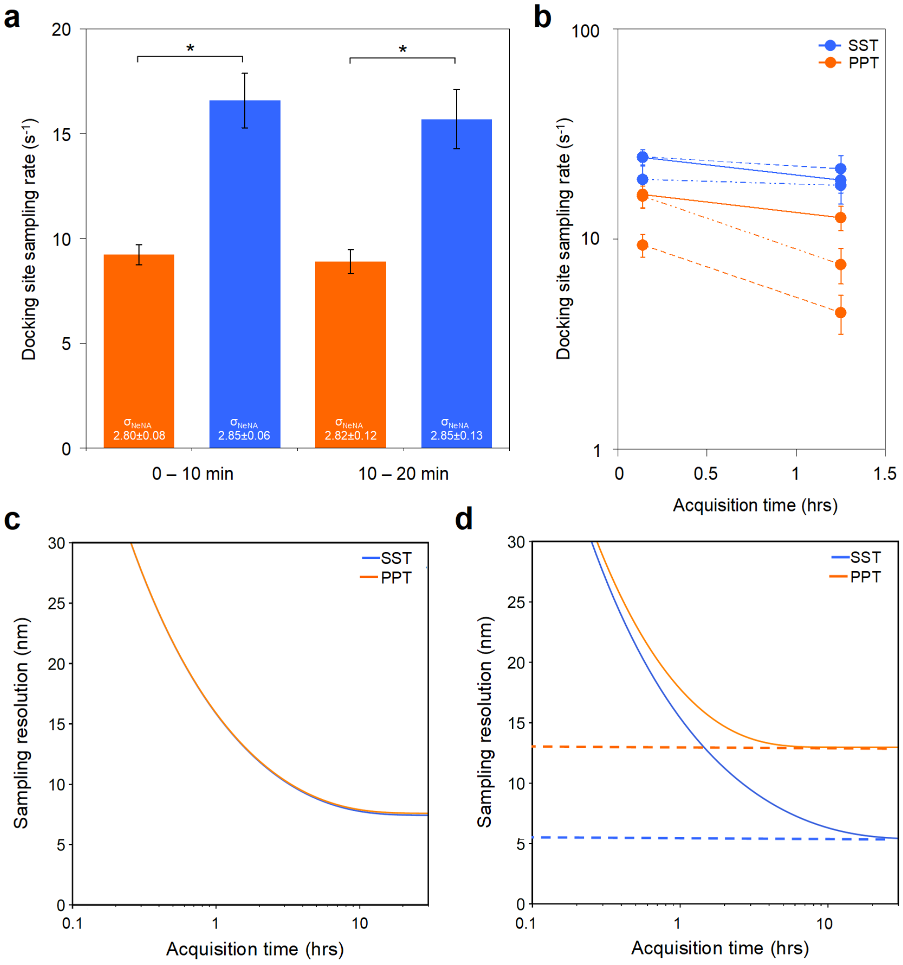


**Fig. S4 | Docking site sampling rate and sampling resolution in SST and PPT.** **a**, Average docking site sampling rate ($s^{-1}$) measured across three wells imaged for 5,000 frames, with analysis performed in two segments of 2,500 frames each. Results compare PPT (orange) and SST (blue). Data are presented as means ±SEM (n=3). * $p < 0.05$, two-tailed unpaired t-test. $\sigma_{NeNA}$ values are presented as means ±SD (n=3). **b**, Docking site sampling rate over three successive 1.5-hour imaging sessions for both PPT (orange) and SST (blue): initial imaging immediately after buffer preparation (solid lines), after 24-hour sample storage (dashed lines), and after one month of storage at room-temperature (dotted lines). Data are presented as means ±SEM (n=11). **c**, Resolution as a function of acquisition time (*T*), estimated damage rates in SST and PPT immediately after preparation. Under these conditions, the achievable sampling resolution is comparable in both systems, plateauing at approximately 7 nm. d, Resolution as a function of acquisition time (T) on logarithmic scale derived from estimated damage rates in SST and PPT buffers, 24 hours after buffer preparation, illustrating how resolution improves through enhanced sampling density. At early time points, the number of accumulated blinks increases linearly; however, over longer durations, docking strand damage limits the number of independent localizations, causing resolution to plateau (dashed lines). SST achieves more than a twofold improvement in resolution compared to PPT.

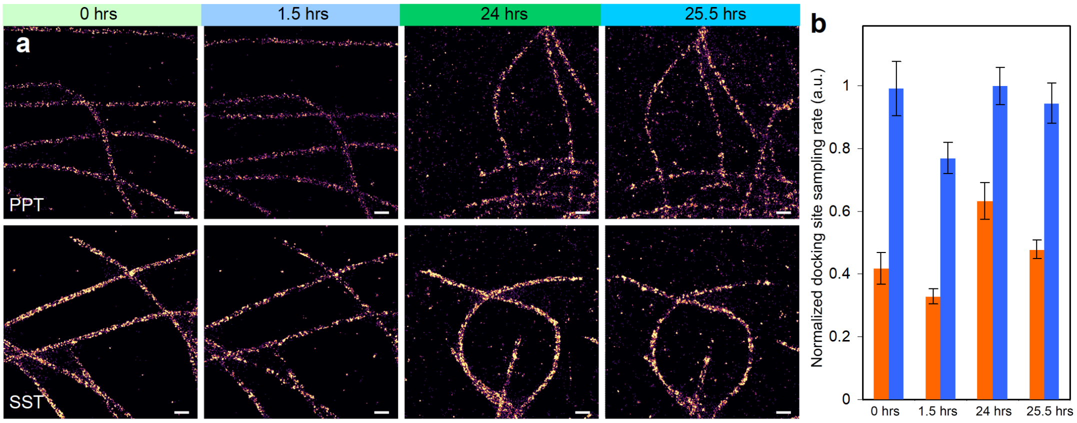


**Fig. S5 | Comparison of microtubule image quality in PPT and SST. a**, Magnified sections highlighting image quality in PPT (top) and SST (bottom) over two 1.5-hour imaging sessions separated by 24 hours of idle time. Scale bars: 250 nm. **b**, Due to the lack of single docking strand resolution, 12 microtubule segments per FOV were selected for analysis, each 4 camera pixels (~520 nm) in length. Docking site sampling rates were quantified for each segment under PPT (orange) and SST (blue) conditions, and normalized to the highest number of localizations observed in any one image. Data are presented as means ±SEM (n=12).

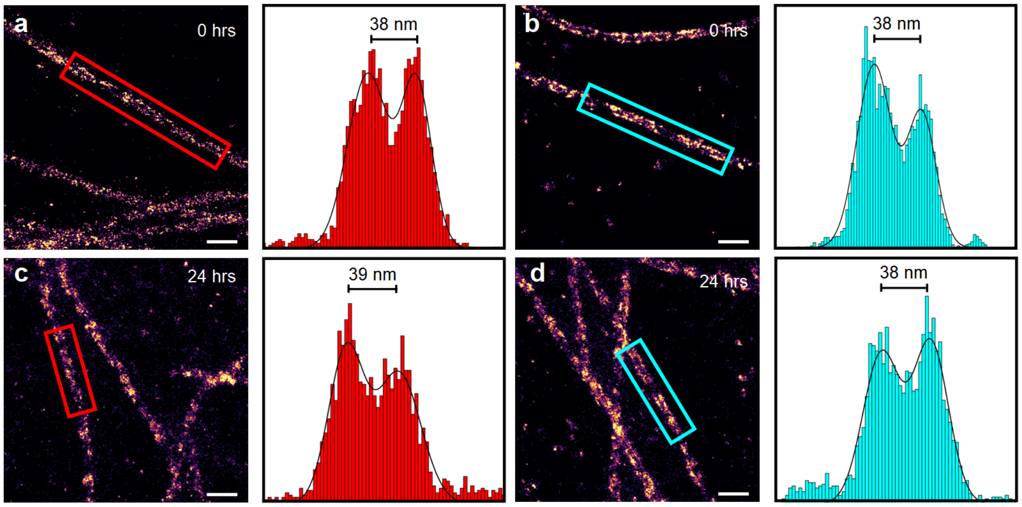


**Fig. S6 | Microtubule cross-sectional fluorescence intensity profiles. a-d,** Microtubule profile analysis comparing PPT (red) and SST (cyan). Left panels: Sections of microtubules with rectangles indicating regions where intensity profiles were extracted. Right panels: Corresponding intensity profiles showing two distinct peaks from labeled microtubule walls projected onto a plane. A double Gaussian fit was applied to determine peak-to-peak distances. Scale bars: 250 nm.

### Supplementary Notes

**Supplementary Note 1.** In order to achieve the highest possible image resolution, DNA-PAINT requires both high per-blink localization precision and high sampling density of distinct events. In practice, DNA-PAINT experiments often last from minutes to many hours to accumulate sufficient sampling density. During these long acquisitions, docking strands can be irreversibly damaged by ROS generated under illumination, reducing the number of docking sites and slowing down further sampling. Photostability thus imposes an ultimate upper limit on resolution, even a system with perfect per-blink precision will fail to resolve fine features if sites are lost before blinking. Localization precision, how accurately the position of a single blink can be estimated, is the equation for the variance of a fitted Gaussian centroid on a pixelated camera[5] and is inversely proportional to the square root of the number of photons, *N*. In DNA-PAINT, however, each docking strand can blink many times, and statistical averaging of *M* independent localizations from the same site reduces uncertainty by a factor of $\sqrt{M}$ . In principle, such averaging can push precision into the sub-nanometer range, provided the camera, optics, and sample remain stable.

While high per-blink precision can, in theory, reach the sub-nanometer regime with sufficient photon counts, practical limitations, including docking strand length, linker flexibility, and biological variability, place additional constraints on achievable resolution. These effects are particularly important when imaging unknown biological structures, where docking site positions are not predetermined. In such cases, achieving high sampling density becomes essential to resolve fine structural details.

However, when imaging an unknown biological structure, one cannot assume knowledge of docking strand positions. If docking sites are arranged in a dense pattern to label a structure of interest, achieving a high distinct localization density across the region becomes critical. The Nyquist-Shannon sampling theorem is a common method for linking image resolution to molecular sampling density[6,7]. In two dimensions, the density of distinct localizations satisfies

$$d_{\text{Nyquist}} = \frac{2}{\sqrt{\rho}} \tag{1}$$

where $\rho$ is the density of localizations per unit area. If each site produces *M* independent localizations, the effective sampling density of localizations in the image is $\rho_{loc} = \rho_0 M$ , where $\rho_0$ is the density of potential binding locations, in other words docking strands, per unit area. This gives

$$d_{\text{Nyquist}} = \frac{2}{\sqrt{\rho_0 M}} \tag{2}$$

Thus even an imaging system capable of sub-nanometer photon-limited precision will only be able to resolve features of size $d_{\text{Nyquist}}$ if the sampling density is sufficiently high. A reduction in available docking sites over time directly constrains the highest resolvable spatial frequencies.

In order to estimate how many blinks per site accumulate over a continuous acquisition time, *T*, each docking strand can be described as undergoing three competing first-order processes: binding of an imager at a rate $k_{\text{on}}\left[c\right]$ (units $s^{-1}$, proportional to imager concentration *c*), unbinding at a rate $k_{\text{off}}$ ($s^{-1}$), and irreversible damage at a rate $k_{\text{dmg}}$ ($s^{-1}$)[8]. The only process that reduces site availability is damage. Let *S(t)* denote the probability that a site remains available at time *t*

$$\frac{dS}{dt} = -k_{\text{dmg}} \Rightarrow S\left(t\right) = e^{-k_{\text{dmg}}t} \tag{3}$$

The instantaneous rate of new blink events is $k_{\text{on}}\left[c\right]S\left(t\right)$. Integrating from time zero to *T* yields the total cumulative number of blinks per site:

$$M(t) = \int_0^T k_{\text{on}}[c] e^{-k_{\text{dmg}} t} = \frac{k_{\text{on}}[c]}{k_{\text{dmg}}}\left(1 - e^{-k_{\text{dmg}} t}\right) \quad (4)$$

At short times $\left(k_{\text{dmg}} T \ll 1\right)$, blink count grows linearly as $k_{\text{on}}[c] S(t)$ and at long times $\left(k_{\text{dmg}} T \gg 1\right)$, blink accumulation saturates at $k_{\text{on}}[c] / k_{\text{dmg}}$, due to the progressive loss of active docking sites through irreversible damage.

The damage rate $k_{\text{dmg}}$, defines the characteristic half-life for docking strand survival as

$$t_{1/2} = \frac{\ln 2}{k_{\text{dmg}}} \quad (5)$$

We developed a simple predictive model using damage rates estimated from the known docking site positions in the 20 nm grid origami. Immediately after buffer preparation, both PPT and SST produced comparable damage rates and predicted sampling-limited resolution of approximately 7 nm (Fig. S4c). However, this estimate is based on a 1.5-hour acquisition prior to PPT degradation and likely overstates resolution achievable in longer experiments. To reflect more realistic usage, we focused our analysis on the 24-hour sample storage condition, which offers a clearer comparison of long-term OSS performance.

Based on our measurements of docking site sampling rates, we estimated damage rates of approximately $2.1\times10^{-4}$ $s^{-1}$ (half-life ~55 minutes) for PPT, and a significantly lower rate of $3.6\times10^{-5}$ $s^{-1}$ (half-life ~5.4 hours) for SST. Using these two regimes, we modeled the evolution of sampling resolution over a 24-hour period.

To calculate Nyquist-limited resolution over any acquisition duration, we substitute Eq. (4) into Eq. (2), yielding the final expression for the estimated sampling resolution:

$$d_{\text{Nyquist}}(T) = 2\sqrt{\frac{k_{\text{dmg}}}{\rho_0 k_{\text{on}}[c]\left(1 - e^{-k_{\text{dmg}} t}\right)}} \quad (6)$$

In PPT, resolution gains diminish after about one hour and plateau at approximately 13 nm. In contrast, SST can, in principle, achieve resolutions below 6 nm with multi-hour acquisitions (Fig. S4d). Overall, DNA-PAINT image acquisitions using SST are expected to achieve more than a twofold improvement in resolution when imaging durations are sufficiently extended.

### Supplementary Tables

**Supplementary Table 1 | List of core staples.** The core origami staple strand sequences were exported in 96-well plate format using Picasso Design[1] and ordered as 96-well plates from IDT at 100 nmole with standard desalting and adjusted to 100mM in IDTE buffer. The empty positions contain place holders for biotinylated staple oligos which were ordered separately (sequences of biotinylated oligos can be found in **Supplementary Table 2**). Note that the first number indicates the plate number (i.e. 1A1 = position A1 on plate 1).

| Plate/ Position | Name | Sequence |
|---|---|---|
| **Plate 1** | | |
| 1A1 | 21[32]23[31]BLK | TTTTCACTCAAAGGGCGAAAAACCATCACC |
| 1A2 | 19[32]21[31]BLK | GTCGACTTCGGCCAACGCGCGGGGTTTTTC |
| 1A3 | 17[32]19[31]BLK | TGCATCTTTCCCAGTCACGACGGCCTGCAG |
| 1A4 | 15[32]17[31]BLK | TAATCAGCGGATTGACCGTAATCGTAACCG |
| 1A5 | 13[32]15[31]BLK | AACGCAAAATCGATGAACGGTACCGGTTGA |
| 1A6 | 11[32]13[31]BLK | AACAGTTTTGTACCAAAAACATTTTATTTC |
| 1A7 | 9[32]11[31]BLK | TTTACCCCAACATGTTTTAAATTTCCATAT |
| 1A8 | 7[32]9[31]BLK | TTTAGGACAAATGCTTTAAACAATCAGGTC |
| 1A9 | 5[32]7[31]BLK | CATCAAGTAAAACGAACTAACGAGTTGAGA |
| 1A10 | 3[32]5[31]BLK | AATACGTTTGAAAGAGGACAGACTGACCTT |
| 1A11 | 1[32]3[31]BLK | AGGCTCCAGAGGCTTTGAGGACACGGGTAA |
| 1A12 | 0[47]1[31]BLK | AGAAAGGAACAACTAAAGGAATTCAAAAAAA |
| 1B1 | 23[32]22[48]BLK | CAAATCAAGTTTTTTGGGGTCGAAACGTGGA |
| 1B2 | 22[47]20[48]BLK | CTCCAACGCAGTGAGACGGGCAACCAGCTGCA |
| 1B3 | 20[47]18[48]BLK | TTAATGAACTAGAGGATCCCCGGGGGGTAACG |
| 1B4 | 18[47]16[48]BLK | CCAGGGTTGCCAGTTTGAGGGGACCCGTGGGA |
| 1B5 | 16[47]14[48]BLK | ACAAACGGAAAAGCCCCAAAAACACTGGAGCA |
| 1B6 | 14[47]12[48]BLK | AACAAGAGGGATAAAAATTTTTAGCATAAAGC |
| 1B7 | 12[47]10[48]BLK | TAAATCGGGATTCCCAATTCTGCGATATAATG |
| 1B8 | 10[47]8[48]BLK | CTGTAGCTTGACTATTATAGTCAGTTCATTGA |
| 1B9 | 8[47]6[48]BLK | ATCCCCCTATACCACATTCAACTAGAAAAATC |
| 1B10 | 6[47]4[48]BLK | TACGTTAAAGTAATCTTGACAAGAACCGAACT |
| 1B11 | 4[47]2[48]BLK | GACCAACTAATGCCACTACGAAGGGGGTAGCA |
| 1B12 | 2[47]0[48]BLK | ACGGCTACAAAAGGAGCCTTTAATGTGAGAAT |
| 1C1 | 21[56]23[63]BLK | AGCTGATTGCCCTTCAGAGTCCACTATTAAAGGGTGCCGT |
| 1C2 | | |
| 1C3 | | |
| 1C4 | 15[64]18[64]BLK | GTATAAGCCAACCCGTCGGATTCTGACGACAGTATCGGCCGCAAGGCG |

| | | |
|---|---|---|
| 1C5 | 13[64]15[63]BLK | TATATTTTGTCATTGCCTGAGAGTGGAAGATT |
| 1C6 | 11[64]13[63]BLK | GATTTAGTCAATAAAGCCTCAGAGAACCCTCA |
| 1C7 | 9[64]11[63]BLK | CGGATTGCAGAGCTTAATTGCTGAAACGAGTA |
| 1C8 | 7[56]9[63]BLK | ATGCAGATACATAACGGGAATCGTCATAAATAAAGCAAAG |
| 1C9 | | |
| 1C10 | | |
| 1C11 | 1[64]4[64]BLK | TTTATCAGGACAGCATCGGAACGACACCAACCTAAAACGAGGTCAATC |
| 1C12 | 0[79]1[63]BLK | ACAACTTTCAACAGTTTCAGCGGATGTATCGG |
| 1D1 | 23[64]22[80]BLK | AAAGCACTAAATCGGAACCCTAATCCAGTT |
| 1D2 | 22[79]20[80]BLK | TGGAACAACCGCCTGGCCCTGAGGCCCGCT |
| 1D3 | 20[79]18[80]BLK | TTCCAGTCGTAATCATGGTCATAAAAGGGG |
| 1D4 | 18[79]16[80]BLK | GATGTGCTTCAGGAAGATCGCACAATGTGA |
| 1D5 | 16[79]14[80]BLK | GCGAGTAAAAATATTTAAATTGTTACAAAG |
| 1D6 | 14[79]12[80]BLK | GCTATCAGAAATGCAATGCCTGAATTAGCA |
| 1D7 | 12[79]10[80]BLK | AAATTAAGTTGACCATTAGATACTTTTGCG |
| 1D8 | 10[79]8[80]BLK | GATGGCTTATCAAAAAGATTAAGAGCGTCC |
| 1D9 | 8[79]6[80]BLK | AATACTGCCCAAAAGGAATTACGTGGCTCA |
| 1D10 | 6[79]4[80]BLK | TTATACCACCAAATCAACGTAACGAACGAG |
| 1D11 | 4[79]2[80]BLK | GCGCAGACAAGAGGCAAAAGAATCCCTCAG |
| 1D12 | 2[79]0[80]BLK | CAGCGAAACTTGCTTTCGAGGTGTTGCTAA |
| 1E1 | 21[96]23[95]BLK | AGCAAGCGTAGGGTTGAGTGTTGTAGGGAGCC |
| 1E2 | 19[96]21[95]BLK | CTGTGTGATTGCGTTGCGCTCACTAGAGTTGC |
| 1E3 | 17[96]19[95]BLK | GCTTTCCGATTACGCCAGCTGGCGGCTGTTTC |
| 1E4 | 15[96]17[95]BLK | ATATTTTGGCTTTCATCAACATTATCCAGCCA |
| 1E5 | 13[96]15[95]BLK | TAGGTAAACTATTTTTGAGAGATCAAACGTTA |
| 1E6 | 11[96]13[95]BLK | AATGGTCAACAGGCAAGGCAAAGAGTAATGTG |
| 1E7 | 9[96]11[95]BLK | CGAAAGACTTTGATAAGAGGTCATATTTCGCA |
| 1E8 | 7[96]9[95]BLK | TAAGAGCAAATGTTTAGACTGGATAGGAAGCC |
| 1E9 | 5[96]7[95]BLK | TCATTCAGATGCGATTTTAAGAACAGGCATAG |
| 1E10 | 3[96]5[95]BLK | ACACTCATCCATGTTACTTAGCCGAAAGCTGC |
| 1E11 | 1[96]3[95]BLK | AAACAGCTTTTTGCGGGATCGTCAACACTAAA |
| 1E12 | 0[111]1[95]BLK | TAAATGAATTTTCTGTATGGGATTAATTTCTT |
| 1F1 | 23[96]22[112]BLK | CCCGATTTAGAGCTTGACGGGGAAAAAGAATA |
| 1F2 | 22[111]20[112]BLK | GCCCGAGAGTCCACGCTGGTTTGCAGCTAACT |
| 1F3 | 20[111]18[112]BLK | CACATTAAAATTGTTATCCGCTCATGCGGGCC |
| 1F4 | 18[111]16[112]BLK | TCTTCGCTGCACCGCTTCTGGTGCGGCCTTCC |
| 1F5 | 16[111]14[112]BLK | TGTAGCCATTAAAATTCGCATTAAATGCCGGA |

| | | |
|---|---|---|
| 1F6 | 14[111]12[112]BLK | GAGGGTAGGATTCAAAAGGGTGAGACATCCAA |
| 1F7 | 12[111]10[112]BLK | TAAATCATATAACCTGTTTAGCTAACCTTTAA |
| 1F8 | 10[111]8[112]BLK | TTGCTCCTTTCAAATATCGCGTTTGAGGGGGT |
| 1F9 | 8[111]6[112]BLK | AATAGTAAACACTATCATAACCCTCATTGTGA |
| 1F10 | 6[111]4[112]BLK | ATTACCTTTGAATAAGGCTTGCCCAAATCCGC |
| 1F11 | 4[111]2[112]BLK | GACCTGCTCTTTGACCCCCAGCGAGGGAGTTA |
| 1F12 | 2[111]0[112]BLK | AAGGCCGCTGATACCGATAGTTGCGACGTTAG |
| 1G1 | 21[120]23[127]BLK | CCCAGCAGGCGAAAAATCCCTTATAAATCAAGCCGGCG |
| 1G2 | | |
| 1G3 | | |
| 1G4 | 15[128]18[128]BLK | TAAATCAAAATAATTCGCGTCTCGGAAACCAGGCAAAGGGAAGG |
| 1G5 | 13[128]15[127]BLK | GAGACAGCTAGCTGATAAATTAATTTTTGT |
| 1G6 | 11[128]13[127]BLK | TTTGGGGATAGTAGTAGCATTAAAAGGCCG |
| 1G7 | 9[128]11[127]BLK | GCTTCAATCAGGATTAGAGAGTTATTTTCA |
| 1G8 | 7[120]9[127]BLK | CGTTTACCAGACGACAAAGAAGTTTTGCCATAATTCGA |
| 1G9 | | |
| 1G10 | | |
| 1G11 | 1[128]4[128]BLK | TGACAACTCGCTGAGGCTTGCATTATACCAAGCGCGATGATAAA |
| 1G12 | 0[143]1[127]BLK | TCTAAAGTTTTGTCGTCTTTCCAGCCGACAA |
| 1H1 | 21[160]22[144]BLK | TCAATATCGAACCTCAAATATCAATTCCGAAA |
| 1H2 | 19[160]20[144]BLK | GCAATTCACATATTCCTGATTATCAAAGTGTA |
| 1H3 | 17[160]18[144]BLK | AGAAAACAAAGAAGATGATGAAACAGGCTGCG |
| 1H4 | 15[160]16[144]BLK | ATCGCAAGTATGTAAATGCTGATGATAGGAAC |
| 1H5 | 13[160]14[144]BLK | GTAATAAGTTAGGCAGAGGCATTTATGATATT |
| 1H6 | 11[160]12[144]BLK | CCAATAGCTCATCGTAGGAATCATGGCATCAA |
| 1H7 | 9[160]10[144]BLK | AGAGAGAAAAAAATGAAAATAGCAAGCAAACT |
| 1H8 | 7[160]8[144]BLK | TTATTACGAAGAACTGGCATGATTGCGAGAGG |
| 1H9 | 5[160]6[144]BLK | GCAAGGCCTCACCAGTAGCACCATGGGCTTGA |
| 1H10 | 3[160]4[144]BLK | TTGACAGGCCACCACCAGAGCCGCGATTTGTA |
| 1H11 | 1[160]2[144]BLK | TTAGGATTGGCTGAGACTCCTCAATAACCGAT |
| 1H12 | 0[175]0[144]BLK | TCCACAGACAGCCCTCATAGTTAGCGTAACGA |
| **Plate 2** | | |
| 2A1 | 23[128]23[159]BLK | AACGTGGCGAGAAAGGAAGGGAAACCAGTAA |
| 2A2 | 22[143]21[159]BLK | TCGGCAAATCCTGTTTGATGGTGGACCCTCAA |
| 2A3 | 20[143]19[159]BLK | AAGCCTGGTACGAGCCGGAAGCATAGATGATG |
| 2A4 | 18[143]17[159]BLK | CAACTGTTGCGCCATTCGCCATTCAAACATCA |
| 2A5 | 16[143]15[159]BLK | GCCATCAAGCTCATTTTTTAACCACAAATCCA |

| | | |
|---|---|---|
| 2A6 | 14[143]13[159]BLK | CAACCGTTTCAAATCACCATCAATTCGAGCCA |
| 2A7 | 12[143]11[159]BLK | TTCTACTACGCGAGCTGAAAAGGTTACCGCGC |
| 2A8 | 10[143]9[159]BLK | CCAACAGGAGCGAACCAGACCGGAGCCTTTAC |
| 2A9 | 8[143]7[159]BLK | CTTTTGCAGATAAAAACCAAAATAAAGACTCC |
| 2A10 | 6[143]5[159]BLK | GATGGTTTGAACGAGTAGTAAATTTACCATTA |
| 2A11 | 4[143]3[159]BLK | TCATCGCCAACAAAGTACAACGGACGCCAGCA |
| 2A12 | 2[143]1[159]BLK | ATATTCGGAACCATCGCCCACGCAGAGAAGGA |
| 2B1 | 23[160]22[176]BLK | TAAAAGGGACATTCTGGCCAACAAAGCATC |
| 2B2 | 22[175]20[176]BLK | ACCTTGCTTGGTCAGTTGGCAAAGAGCGGA |
| 2B3 | 20[175]18[176]BLK | ATTATCATTCAATATAATCCTGACAATTAC |
| 2B4 | 18[175]16[176]BLK | CTGAGCAAAAATTAATTACATTTTGGGTTA |
| 2B5 | 16[175]14[176]BLK | TATAACTAACAAAGAACGCGAGAACGCCAA |
| 2B6 | 14[175]12[176]BLK | CATGTAATAGAATATAAAGTACCAAGCCGT |
| 2B7 | 12[175]10[176]BLK | TTTTATTTAAGCAAATCAGATATTTTTTGT |
| 2B8 | 10[175]8[176]BLK | TTAACGTCTAACATAAAAACAGGTAACGGA |
| 2B9 | 8[175]6[176]BLK | ATACCCAACAGTATGTTAGCAAATTAGAGC |
| 2B10 | 6[175]4[176]BLK | CAGCAAAAGGAAACGTCACCAATGAGCCGC |
| 2B11 | 4[175]2[176]BLK | CACCAGAAAGGTTGAGGCAGGTCATGAAAG |
| 2B12 | 2[175]0[176]BLK | TATTAAGAAGCGGGGTTTTGCTCGTAGCAT |
| 2C1 | 21[184]23[191]BLK | TCAACAGTTGAAAGGAGCAAATGAAAAATCTAGAGATAGA |
| 2C2 | | |
| 2C3 | | |
| 2C4 | 15[192]18[192]BLK | TCAAATATAACCTCCGGCTTAGGTAACAATTTCATTTGAAGGCGAATT |
| 2C5 | 13[192]15[191]BLK | GTAAAGTAATCGCCATATTTAACAAAACTTTT |
| 2C6 | 11[192]13[191]BLK | TATCCGGTCTCATCGAGAACAAGCGACAAAAG |
| 2C7 | 9[192]11[191]BLK | TTAGACGGCCAAATAAGAAACGATAGAAGGCT |
| 2C8 | 7[184]9[191]BLK | CGTAGAAAATACATACCGAGGAAACGCAATAAGAAGCGCA |
| 2C9 | | |
| 2C10 | | |
| 2C11 | 1[192]4[192]BLK | GCGGATAACCTATTATTCTGAAACAGACGATTGGCCTTGAAGAGCCAC |
| 2C12 | 0[207]1[191]BLK | TCACCAGTACAAACTACAACGCCTAGTACCAG |
| 2D1 | 23[192]22[208]BLK | ACCCTTCTGACCTGAAAGCGTAAGACGCTGAG |
| 2D2 | 22[207]20[208]BLK | AGCCAGCAATTGAGGAAGGTTATCATCATTTT |
| 2D3 | 20[207]18[208]BLK | GCGGAACATCTGAATAATGGAAGGTACAAAAT |
| 2D4 | 18[207]16[208]BLK | CGCGCAGATTACCTTTTTTAATGGGAGAGACT |
| 2D5 | 16[207]14[208]BLK | ACCTTTTTATTTTAGTTAATTTCATAGGGCTT |
| 2D6 | 14[207]12[208]BLK | AATTGAGAATTCTGTCCAGACGACTAAACCAA |

| | | |
|---|---|---|
| 2D7 | 12[207]10[208]BLK | GTACCGCAATTCTAAGAACGCGAGTATTATTT |
| 2D8 | 10[207]8[208]BLK | ATCCCAATGAGAATTAACTGAACAGTTACCAG |
| 2D9 | 8[207]6[208]BLK | AAGGAAACATAAAGGTGGCAACATTATCACCG |
| 2D10 | 6[207]4[208]BLK | TCACCGACGCACCGTAATCAGTAGCAGAACCG |
| 2D11 | 4[207]2[208]BLK | CCACCCTCTATTCACAAACAAATACCTGCCTA |
| 2D12 | 2[207]0[208]BLK | TTTCGGAAGTGCCGTCGAGAGGGTGAGTTTCG |
| 2E1 | 21[224]23[223]BLK | CTTTAGGGCCTGCAACAGTGCCAATACGTG |
| 2E2 | 19[224]21[223]BLK | CTACCATAGTTTGAGTAACATTTAAAATAT |
| 2E3 | 17[224]19[223]BLK | CATAAATCTTTGAATACCAAGTGTTAGAAC |
| 2E4 | 15[224]17[223]BLK | CCTAAATCAAAATCATAGGTCTAAACAGTA |
| 2E5 | 13[224]15[223]BLK | ACAACATGCCAACGCTCAACAGTCTTCTGA |
| 2E6 | 11[224]13[223]BLK | GCGAACCTCCAAGAACGGGTATGACAATAA |
| 2E7 | 9[224]11[223]BLK | AAAGTCACAAAATAAACAGCCAGCGTTTTA |
| 2E8 | 7[224]9[223]BLK | AACGCAAAGATAGCCGAACAAACCCTGAAC |
| 2E9 | 5[224]7[223]BLK | TCAAGTTTCATTAAAGGTGAATATAAAAGA |
| 2E10 | 3[224]5[223]BLK | TTAAAGCCAGAGCCGCCACCCTCGACAGAA |
| 2E11 | 1[224]3[223]BLK | GTATAGCAAACAGTTAATGCCCAATCCTCA |
| 2E12 | 0[239]1[223]BLK | AGGAACCCATGTACCGTAACACTTGATATAA |
| 2F1 | 23[224]22[240]BLK | GCACAGACAATATTTTTGAATGGGGTCAGTA |
| 2F2 | 22[239]20[240]BLK | TTAACACCAGCACTAACAACTAATCGTTATTA |
| 2F3 | 20[239]18[240]BLK | ATTTTAAAATCAAAATTATTTGCACGGATTCG |
| 2F4 | 18[239]16[240]BLK | CCTGATTGCAATATATGTGAGTGATCAATAGT |
| 2F5 | 16[239]14[240]BLK | GAATTTATTTAATGGTTTGAAATATTCTTACC |
| 2F6 | 14[239]12[240]BLK | AGTATAAAGTTCAGCTAATGCAGATGTCTTTC |
| 2F7 | 12[239]10[240]BLK | CTTATCATTCCCGACTTGCGGGAGCCTAATTT |
| 2F8 | 10[239]8[240]BLK | GCCAGTTAGAGGGTAATTGAGCGCTTTAAGAA |
| 2F9 | 8[239]6[240]BLK | AAGTAAGCAGACACCACGGAATAATATTGACG |
| 2F10 | 6[239]4[240]BLK | GAAATTATTGCCTTTAGCGTCAGACCGGAACC |
| 2F11 | 4[239]2[240]BLK | GCCTCCCTCAGAATGGAAAGCGCAGTAACAGT |
| 2F12 | 2[239]0[240]BLK | GCCCGTATCCGGAATAGGTGTATCAGCCCAAT |
| 2G1 | 21[248]23[255]BLK | AGATTAGAGCCGTCAAAAAACAGAGGTGAGGCCTATTAGT |
| 2G2 | | |
| 2G3 | | |
| 2G4 | 15[256]18[256]BLK | GTGATAAAAAGACGCTGAGAAGAGATAACCTTGCTTCTGTTCGGGAGA |
| 2G5 | 13[256]15[255]BLK | GTTTATCAATATGCGTTATACAAACCGACCGT |
| 2G6 | 11[256]13[255]BLK | GCCTTAAACCAATCAATAATCGGCACGCGCCT |
| 2G7 | 9[256]11[255]BLK | GAGAGATAGAGCGTCTTTCCAGAGGTTTTGAA |

| | | |
|---|---|---|
| 2G8 | 7[248]9[255]BLK | GTTTATTTTGTCACAATCTTACCGAAGCCCTTTAATATCA |
| 2G9 | | |
| 2G10 | | |
| 2G11 | 1[256]4[256]BLK | CAGGAGGTGGGGTCAGTGCCTTGAGTCTCTGAATTTACCGGGAACCAG |
| 2G12 | 0[271]1[255]BLK | CCACCCTCATTTTCAGGGATAGCAACCGTACT |
| 2H1 | 23[256]22[272]BLK | CTTTAATGCGCGAACTGATAGCCCCACCAG |
| 2H2 | 22[271]20[272]BLK | CAGAAGATTAGATAATACATTTGTCGACAA |
| 2H3 | 20[271]18[272]BLK | CTCGTATTAGAAATTGCGTAGATACAGTAC |
| 2H4 | 18[271]16[272]BLK | CTTTTACAAAATCGTCGCTATTAGCGATAG |
| 2H5 | 16[271]14[272]BLK | CTTAGATTTAAGGCGTTAAATAAAGCCTGT |
| 2H6 | 14[271]12[272]BLK | TTAGTATCACAATAGATAAGTCCACGAGCA |
| 2H7 | 12[271]10[272]BLK | TGTAGAAATCAAGATTAGTTGCTCTTACCA |
| 2H8 | 10[271]8[272]BLK | ACGCTAACACCCACAAGAATTGAAAATAGC |
| 2H9 | 8[271]6[272]BLK | AATAGCTATCAATAGAAAATTCAACATTCA |
| 2H10 | 6[271]4[272]BLK | ACCGATTGTCGGCATTTTCGGTCATAATCA |
| 2H11 | 4[271]2[272]BLK | AAATCACCTTCCAGTAAGCGTCAGTAATAA |
| 2H12 | 2[271]0[272]BLK | GTTTTAACTTAGTACCGCCACCCAGAGCCA |

**Supplementary Table 2 | List of biotinylated staples**. Biotinilated staples were ordered from IDT at 100 nmole with standard desalting and adjusted to 100mM in IDTE buffer.

| No | Name | Sequence | Mod |
|---|---|---|---|
| 1 | 18[63]20[56]BIOTIN | ATTAAGTTTACCGAGCTCGAATTCGGGAAACCTGTCGTGC | 5'-BT |
| 2 | 4[63]6[56]BIOTIN | ATAAGGGAACCGGATATTCATTACGTCAGGACGTTGGGAA | 5'-BT |
| 3 | 18[127]20[120]BIOTIN | GCGATCGGCAATTCCACACAACAGGTGCCTAATGAGTG | 5'-BT |
| 4 | 4[127]6[120]BIOTIN | TTGTGTCGTGACGAGAAACACCAAATTTCAACTTTAAT | 5'-BT |
| 5 | 18[191]20[184]BIOTIN | ATTCATTTTTGTTTGGATTATACTAAGAAACCACCAGAAG | 5'-BT |
| 6 | 4[191]6[184]BIOTIN | CACCCTCAGAAACCATCGATAGCATTGAGCCATTTGGGAA | 5'-BT |
| 7 | 18[255]20[248]BIOTIN | AACAATAACGTAAAACAGAAATAAAAATCCTTTGCCCGAA | 5'-BT |
| 8 | 4[255]6[248]BIOTIN | AGCCACCACTGTAGCGCGTTTTCAAGGGAGGGAAGGTAAA | 5'-BT |

**Supplementary Table 3 | Functional staple sequences.** The following functional oligos were used to replace core staple position in the respective DNA origami design ('SD' – fixed single-dye origami[3] and '20nm' – 20-nm grid with 3x4 pattern of docking strands for DNA-PAINT). The docking strand-extended staples were ordered from IDT at 100nmole with standard desalting at 100mM in IDTE buffer for standard docking strands and the fixed-Cy3b modified staple at 250nmole with HPLC purification and lyophilized.

| **Shortname (docking site length)** | **Origami ID** | **Docking strand sequence** | **Concatenated to following staple positions** |
|---|---|---|---|
| 5xCTC | 20nm | TT CTCCTCCTCCTCCTC | 1B3, 2B3, 1F3, 2F3<br>1B7, 2B7, 1F7, 2F7<br>1B11, 2B11, 1F11, 2F11 |
| 4T_Cy3b | SD | TTTT-Cy3b | 2B7 |

**Supplementary Table 4 | Imager strand sequence.** Cy3b-labeled imager strand was ordered from IDT at 250nmole with HPLC purification and lyophilized. The stock solution was adjusted to 100µM and stored at -20 °C. Working aliquots at 1 µM were stored in the dark at 4 °C.

| **Shortname (docking site length)** | **Imager sequence** |
|---|---|
| 5xCTC | GAGGAGG-Cy3b |
